# Layer-dependent multiplicative effects of spatial attention on contrast responses in human early visual cortex

**DOI:** 10.1101/2020.02.01.926303

**Authors:** Fanhua Guo, Chengwen Liu, Chencan Qian, Zihao Zhang, Kaibao Sun, Danny JJ Wang, Sheng He, Peng Zhang

**Affiliations:** State Key Laboratory of Brain and Cognitive Science, Institute of Biophysics, Chinese Academy of Sciences, Beijing, 100101, China; University of Chinese Academy of Sciences, Beijing 100049, China; School of Ophthalmology and Optometry and Eye hospital, and State Key Laboratory of Ophthalmology, Optometry and Vision Science, Wenzhou Medical University, Wenzhou, Zhejiang, 325000, China; Stevens Neuroimaging and Informatics Institute, University of Southern California, Los Angeles, CA, United States; Department of Psychology, University of Minnesota, Minneapolis, MN 55455

**Keywords:** Layer, 7T fMRI, Feedforward, Feedback, Spatial attention, Contrast response function

## Abstract

Attention mechanisms at different cortical layers of human visual cortex remain poorly understood. Using submillimeter-resolution fMRI at 7T, we investigated the effects of top-down spatial attention on the contrast responses across different cortical depths in human early visual cortex. Gradient echo (GE) T2* weighted BOLD signal showed an additive effect of attention on contrast responses across cortical depths. Compared to the middle cortical depth, attention modulation was stronger in the superficial and deep depths of V1, and also stronger in the superficial depth of V2 and V3. Using ultra-high resolution (0.3mm in-plane) balanced steady-state free precession (bSSFP) fMRI, a multiplicative scaling effect of attention was found in the superficial and deep layers, but not in the middle layer of V1. Attention modulation of low contrast response was strongest in the middle cortical depths, indicating baseline enhancement or contrast gain of attention modulation on feedforward input. Finally, the additive effect of attention on T2* BOLD can be explained by strong nonlinearity of BOLD signals from large blood vessels, suggesting multiplicative effect of attention on neural activity. These findings support that top-down spatial attention mainly operates through feedback connections from higher order cortical areas, and a distinct mechanism of attention may also be associated with feedforward input through subcortical pathway.

**Highlights:**

- Response or activity gain of spatial attention in superficial and deep layers
- Contrast gain or baseline shift of attention in V1 middle layer
- Nonlinearity of large blood vessel causes additive effect of attention on T2* BOLD

## Introduction

Our eyes are bombarded with an overwhelming amount of visual information. Given limited information processing capacity of our brain, attention is important to selectively process relevant visual information for efficient allocation of computational resources. Directing attention to a spatial location can improve sensitivity and performance of visual tasks at the attended location, by enhancing neural activity throughout the visual hierarchy. There are multiple cortical and subcortical pathways through which attention could possibly act through and modulate early visual activity. However, the neural mechanisms of attention associated with these feedforward and feedback neural circuitries remain poorly understood. Human visual cortex consists of six layers of neurons, with distinct roles in feedforward, feedback and intracortical connections (Felleman and Van Essen 1991). Investigating layer-specific modulation of attention in human visual cortex is critical to understand whether and how attention modulates neural activity through feedforward or feedback visual pathways.

Figure 1A shows the possible feedforward and feedback pathways that could be associated with attention modulation. One possibility is that attention modulates neural activity through an ascending feedforward pathway: SC → TRN → LGN → V1. This hypothesis is supported by electrophysiological evidence showing early attentional enhancement of LGN activity (26 ms and 37 ms in M and P layers, respectively) following suppressed neural activity of thalamic reticular nucleus (TRN, 22 ms) (McAlonan, Cavanaugh et al. 2008), and by tracing studies showing projection to TRN from the deep layer of superior colliculus (SC), an attention control nucleus in the brain stem (Kolmac and Mitrofanis 1998). Since layer 4 of early visual cortex mainly receives feedforward input, if attention modulates neural activity through the feedforward pathway, the effect of attention modulation should be strongest in the middle cortical depth. Here we focus on input activity because blood oxygen level dependent (BOLD) signals are better correlated with local field potentials (LFPs) which reflects the post-synaptic input rather than the spiking output activity (Logothetis, Pauls et al. 2001). Another possibility is that attention operates through feedback connections along descending visual pathways, such as cortico-cortical feedback targeting the superficial and deeper layers (Felleman and Van Essen 1991), or thalamo-cortical feedback from the pulvinar targeting the superficial layers of early visual cortex (Ogren and Hendrickson 1977, Rezak and Benevento 1979, Shipp 2003). Thus if attention mainly modulates neural activity through feedback connections, we expect to see attention modulation predominates in the superficial and/or deep layers. To test these hypotheses of neural mechanisms of attention in the human brain, we need to non-invasively access layer-specific signal in the human visual cortex.

**Figure 1.**
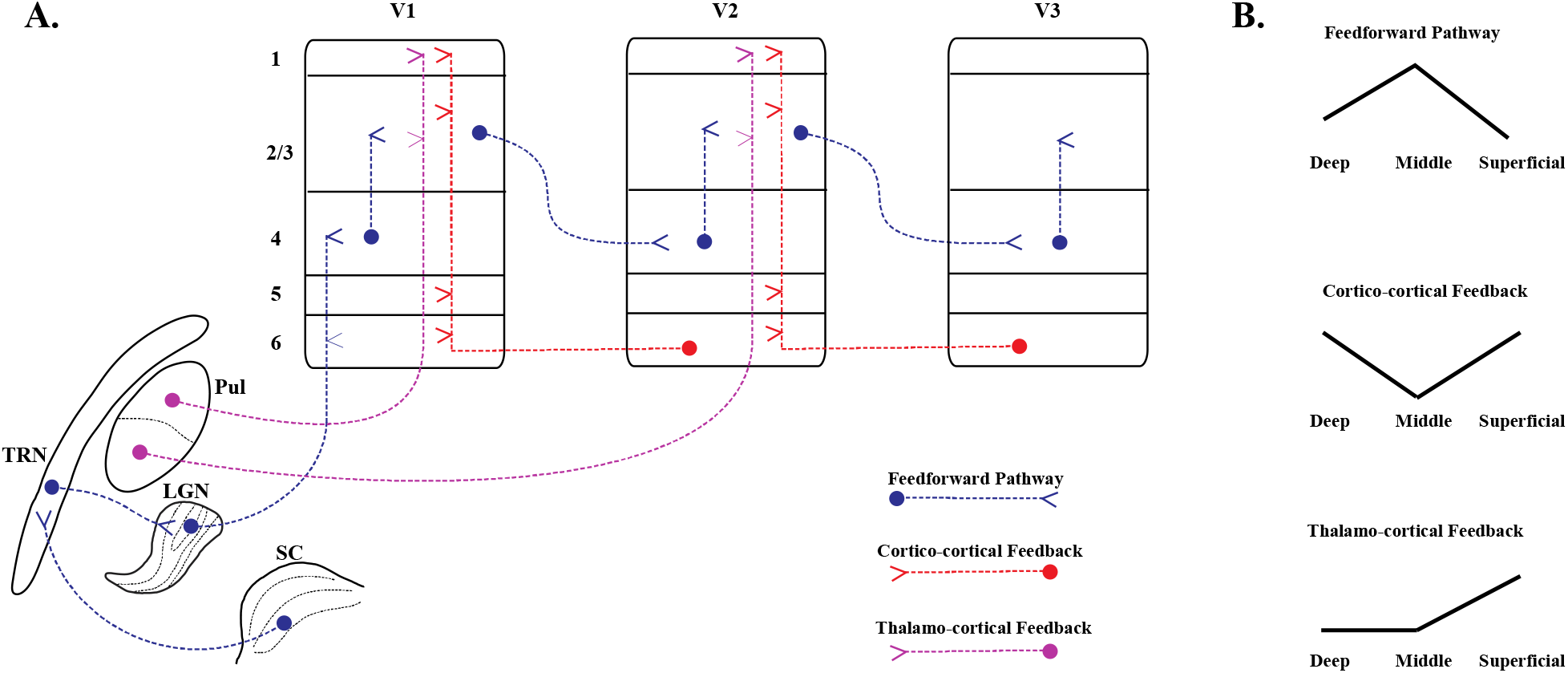
**A.** Possible neural pathways of attention modulation (Felleman and Van Essen 1991, Shipp 2003, McAlonan, Cavanaugh et al. 2008). **B.** Laminar profile of attention modulation in early visual cortex predicted by forward and feedback connections. Abbreviations: SC, superior colliculus, LGN, lateral geniculate nucleus, TRN, thalamic reticular nucleus, Pul, pulvinar.

Recent advances in functional magnetic resonance imaging (fMRI) techniques and data analysis strategies at ultrahigh magnetic field open a window of opportunity to study cortical layer-specific activity non-invasively in the human brain (Polimeni, Renvall et al. 2017, Huber, Uludag et al. 2019). However, current lamina-resolution fMRI techniques mainly used EPI-based sequences, thus one major challenge is image distortion and blur due to B0 inhomogeneity and long EPI readout. An exciting method for minimizing such artifacts is balanced steady-state free precession (bSSFP). Due to extremely short readout time and rapid spin refocusing, bSSFP image has minimal distortion or blur. At short TR, the rapid refocusing mechanism of balanced SSFP suppresses large scale off-resonance effect, and is more sensitive to diffusion activity around small vessels (Lee, Dumoulin et al. 2008, Miller 2012, Scheffler, Heule et al. 2018). Thus the contrast mechanism of rapid bSSFP is similar to T2 weighted spin echo BOLD, and may provide high spatial specificity for lamina-resolution fMRI. Passband bSSFP utilizes the flat portion of off-resonance signal profile to achieve homogeneous and high SNR imaging of a large brain region. These properties make pass-band bSSFP an attractive technique for distortion free high-resolution fMRI at ultra-high field. To the best of our knowledge, pass-band bSSFP fMRI has not yet been applied to study cortical layer-dependent activity in the human brain.

In the current study, we used both traditional GE-EPI (T2* weighted) and passband bSSFP (T2 weighted) fMRI methods at 7T to investigate layer-dependent effect of spatial attention on BOLD contrast response functions. Attention modulation of contrast response functions can provide a quantitative assessment of the gain control mechanisms of attention. Previous behavioral and electrophysiological studies showed that attention modulation on contrast response functions can be horizontal shift (contrast gain) (Reynolds, Pasternak et al. 2000, Martinez-Trujillo and Treue 2002), multiplicative scaling (response or activity gain)(McAdams and Maunsell 1999, Di Russo, Spinelli et al. 2001, Williford and Maunsell 2006), or additive offset (baseline shift) (Kastner, Pinsk et al. 1999, O’Connor, Fukui et al. 2002). These attention effect can be explained by a divisive normalization model of attention (Reynolds and Heeger 2009). However, previous GE BOLD fMRI studies at 3T mainly revealed additive effect of attention on contrast responses (Buracas and Boynton 2007, Li, Lu et al. 2008, Pestilli, Carrasco et al. 2011). Although a number of explanations have been proposed, the answer to the discrepancy between attention effects observed with BOLD fMRI and with electrophysiology remains elusive. One important consideration is that BOLD signal arises from complex vascular responses, which might be differently modulated by attention compared with neural activity. At 3T, T2* BOLD signals are dominated by nonspecific intravascular responses from large draining veins (Zhong, Kennan et al. 1998), which is far from the site of neural activation. At ultra-high magnetic field (7T and above), intravascular response from large draining vein is much reduced (Duong, Yacoub et al. 2003). Higher signal-to-noise ratio at 7T also enables higher spatial resolution to reduce partial volume effect from large pial veins. The refocusing mechanism of T2 weighted fMRI method such as spin echo and balanced SSFP further eliminated the extravascular response from large vessels. Therefore, we expect that 7T fMRI signals with higher tissue specificity (e.g. T2 weighted bSSFP) might help to reveal the neural mechanism attention on contrast responses. Indeed, we found response/activity gain of attention on T2 weighted bSSFP signals in the superficial and deep layers of V1, while attention modulation of low contrast response was strongest in the middle layer. These findings support that top-down spatial attention mainly operates through feedback visual pathways; and at low contrast levels, a distinct mechanism of attention may also be associated with feedforward input through subcortical pathway.

## Methods and Materials

### Participants

Fifteen subjects (6 females, age range 22-39, mean age 26.2) participated in the GE-EPI experiment (Exp 1). Twelve subjects (6 females, age range 22-40, mean age 26.4) participated in the ultra-high resolution bSSFP experiment (Exp 2). All subjects had normal or corrected to normal vision, and gave written informed consent before the experiments. Experimental protocol was approved by institutional review board of the Institute of Biophysics, Chinese Academy of Sciences.

### Stimuli and procedures

Stimuli were generated in MATLAB (Mathworks Inc.) with psychophysics toolbox extension (Brainard 1997, Pelli 1997) on a MacPro computer. Stimuli were presented with a MRI safe projector (1024×768@60Hz) on a translucent screen behind the head coil. Participants viewed the stimuli through a mirror mounted inside the head coil. The color look up table of the projector was calibrated to have a linear luminance output.

Figure 2 shows the stimuli and procedure for experiment 1. Subjects were required to keep fixation during experiment. Before stimulus presentation, a central cue was presented at fixation for 1 second. Then a pair of counter-phase flickering (7.5Hz) square wave checkerboards patterns (0.5 cycles per degree) were presented to the left and right side of fixation at the lower visual field for 16 seconds, followed by 8s of fixation period. The size of the checkerboard discs were 7 degrees in diameter, presented at an eccentricity of 6 degrees. During stimulus presentation, the spatial frequency of the two checkerboard patterns changed 1-3 times randomly and independently. Subjects were asked to pay attention to the cued checkerboard pattern to detect occasional spatial frequency change of the attended stimulus. The checkerboard patterns were presented at five contrast levels (2.5%, 6%, 14.4%, 34.6% and 82.9%). Different contrast conditions were presented in separate runs. The order of different contrast conditions was counterbalanced across subjects. In a pilot experiment before scanning, task difficulties were matched across different contrast conditions with a three-down one-up staircase procedure. Each run lasted 240s, contained 10 stimulus blocks alternating with fixation period. Each subject completed 10 runs of functional scan (two runs for each contrast condition). Stimuli and procedures of experiment 2 were similar as in experiment 1, except as follows. Two wedges of checkerboard stimuli (150 degrees of angles, 0.5 to 5 degrees of eccentricity, 2 cpd spatial frequency, counterphase flickering at 6 Hz) were simultaneously presented to the left and right side of fixation. Stimuli were presented at two contrast levels (5%, 50%). Eight runs of function data were collected for each subject.

**Figure 2.**
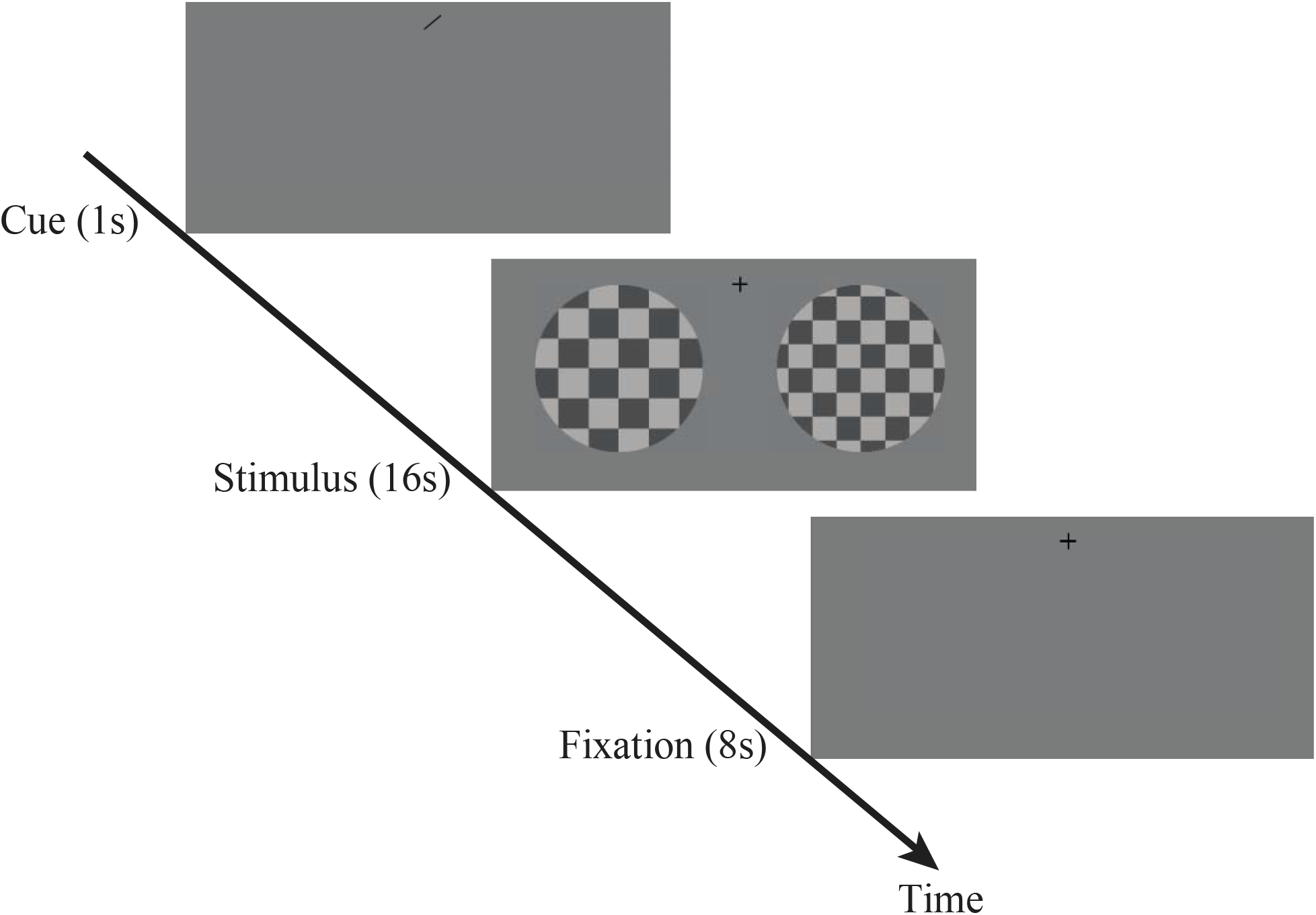
Schematic paradigm and procedure of experiment 1. Before stimulus presentation, an attention cue was presented at the fixation for 1s, then two flickering checkerboard was presented to the left and to the right side of fixation for 16s, followed by 8s fixation period. Subjects’ task was to detect occasional spatial frequency change of the checkerboard stimulus at the cued location.

### MRI data acquisition and analysis for the GE-EPI experiment

#### MRI data acquisition

MRI data were acquired on a 7T MRI scanner (Siemens Magnetom) with a 32-channel receive 1-channel transmit head coil (Nova Medical), from Beijing MRI center for Brain Research (BMCBR). Gradient coil has a maximum amplitude of 70mT/m, 200 us minimum gradient rise time, and 200T/m/s maximum slew rate. In experiment 1, high resolution (0.75 mm isotropic) functional data were collected with a T2*-weighted 2D GE-EPI sequence (37 slices, TR=2667ms, TE=25.6ms, FOV=128*128mm, image matrix = 170*170, GRAPPA acceleration factor = 3, Flip angle=80°, phase encoding direction from Foot to Head). EPI images with reversed phase encoding direction (H to F) were also acquired for distortion correction. Anatomical volumes were acquired with T1, PD and T2* weighted sequences at 0.7 mm isotropic resolution (T1 weighted MPRAGE: 256 sagittal slices, centric phase encoding, acquisition matrix=320*320, Field of view = 223mm*223mm, GRAPPA=3, TR =3000ms, TE=3.76ms,TI = 1250ms, flip angle = 8 °; PD weighted MPRAGE: 256 sagittal slices, matrix = 320*320, Field of view = 223mm*223mm, GRAPPA=3, TR=2340ms, TE=3.38ms, flip angle = 8°; T2* weighted GRE: 208 sagittal slices, matrix = 320*320, Field of view = 223mm*223mm, GRAPPA=3, TR =17ms, TE=12ms, flip angle = 14 °). Respiration and pulse signal were recorded for physiology noise removal.

#### Preprocessing of functional data

Preprocessing of functional data was done in AFNI (cox, 1996), including the following steps: physiological noise removal with retrospective image correction, slice timing correction, EPI image distortion correction with nonlinear warping (PE blip-up), rigid body motion correction, alignment of corrected EPI images to T1-weighted anatomical volume (cost function: lpc), and per run scaling as percent signal change. To minimize image blur, all spatial transformations were combined together and applied to the functional images in one resampling step. General linear models with a fixed HRF (Block4 in AFNI) were used to estimate BOLD signal change from baseline for each stimulus condition. Physiological noise regressors, head motion parameters, as well as their derivatives and square of derivatives, were included as regressors of no interest.

#### Surface segmentation

Proton density images were used to remove spatial intensity inhomogeneities related to transmit-and-receive profile of the RF coils, while also enhancing the contrast between gray and white matter, by dividing T1-weighted by PD-weighted anatomical images (Van de Moortele, Auerbach et al. 2009). The T1/PD anatomical volume was segmented into white matter, gray matter, and CSF using FreeSurfer’s automated procedure with high-resolution option. The results of initial segmentation were visually inspected and manually edited to eliminate dura mater, sinus, etc., ensuring correct surface segmentation (Figure S1). The cortical depth profiles were constructed using an equi-volume model by taking the local curvature of pial and white matter surfaces into account, which has been shown to be more accurate than the equi-distance model (Waehnert, Dinse et al. 2014). We calculated two intermediate surfaces between white matter (WM) and pial surfaces, dividing the gray matter (GM) into three equi-volume layer compartments. For each voxel, we calculated the percentage of volume in the WM, CSF and the three cortical layer compartments. These layer-weights were subsequently used in a spatial regression approach to unmix layer activity (Kok, Bains et al. 2016). This unmixing method helps decreasing the activity correlations across different layers.

#### ROI definition

Retinotopic ROIs of V1-V3 were defined on the cortical surface according to the polar angle atlas from the 7T retinotopic dataset of Human Connectome Project (Benson, Jamison et al. 2018). Only vertices with significant activation (mean response across all conditions, p<0.05 before correction) were included in the analyses. Three approaches were combined to reduce the influence from large pial veins. First, vertices with low EPI intensity from the mean EPI image were removed(Kay, Jamison et al. 2019). The mean EPI image was first bias-corrected to remove non-uniformity of signal intensity (3dUnifize in AFNI), then vertices with an intensity below 75% of the mean intensity of all surface vertices were excluded. Second, vertices with large signal change (3% signal change averaged across all conditions) were excluded (Cheng, Waggoner et al. 2001). Finally, vertices with high intensity value from the PD/T2* image were removed (De Martino, Moerel et al. 2015). Large pial veins has high signal intensity on the PD weighted image whereas low signal intensity on T2* weighted image, thus the ratio between PD and T2* weighted images greatly enhanced the contrast between veins and brain tissue, allowing robust removal of large pial veins.

### MRI data acquisition and analysis for the bSSFP experiment

#### MRI data acquisition

In experiment 2, functional data were acquired with a T2-weighted 2D passband balanced-SSFP sequence (a single 3mm thick slice with 0.3mm in-plane resolution, TE = 2.4ms, TR = 4.8ms, slice acquisition time = 3.2s, FOV = 96mm*96mm, no parallel imaging, Flip angle = 20 deg). Anatomical images were collected with a T1-MP2RAGE sequence (0.7mm isotropic voxels, acquisition matrix = 320*320, GRAPPA = 3, TE = 3.05ms, TR = 4000ms, TI1 = 750 ms, Flip angle = 4 deg, TI2 = 2500 ms, Flip angle = 5 deg, FOV = 224*224 mm^2^). Subjects used bite bars to restrict head motion. To avoid the banding artifact of bSSFP images, we carefully adjusted the RF frequency to keep the region of interest in the middle of the passband of off-resonance profile. A series of bSSFP images were collected with RF frequency shift Δf) from −70Hz to 70Hz with a step of 10Hz (Figure S2). The optimal Δf can be determined from the off-resonance profile from the interested location of V1 gray matter. Three dummy images were acquired to ensure fMRI signal to reach steady-state.

#### MRI data analysis

Motion correction was performed in-plane with three parameters free to change: in-plane rotation and translations (x and y), with the assistance of manual adjustment. T1-MP2RAGE anatomical volume was registered to the mean of motion corrected bSSFP images. Registration results were carefully inspected and adjusted manually if necessary. General linear model with a canonical HRF (Block4 in AFNI) were used to estimate BOLD signal changes from baseline for each stimulus condition. Pial surface was manually defined from the mean of bSSFP image, which provides good contrast between gray matter (GM) and cerebrospinal fluid (CSF). White matter (WM) surface was defined based on T1 contrast between GM and WM from MP2RAGE image, and also referred to the activation map which showed clear boundary at WM surface. For each voxel in the gray matter ROI, cortical depth was calculated by the normalized distance to the pial and WM surface. The laminar profile was subdivided into 20 equal-distance bins.

#### Statistical Analysis

Given the relatively small sample size of our study, we used non-parametric bootstrapping method for the hypothesis tests of cortical depth-dependent effect. For example, to estimate the statistics of model parameters, the entire dataset of each subject was treated as a sampling unit. The group averaged CRFs from the resampled datasets were calculated for each cortical depth of different ROIs and attention conditions. Then parameters from the baseline shift model (R_max_, b, C_50_) were estimated. The bootstrapping procedure was repeated 10,000 times. A null distribution was generated by shifting the mean of bootstrapped distribution to the test value, and then two-sided p value was derived from the null distribution. Cluster-based permutation test was used for multiple comparison corrections for the clustered significance of a range of cortical depths in Figure 6A (middle panel)(Nichols and Holmes 2002). A p value was first calculated for each cortical depth with bootstrapping method. To correct the p value for the continuous range (or cluster) of cortical depths that reached significance, a null distribution was generated for the size of significant cluster. During each permutation, the dataset from each subject was randomly inverted with a minus sign (e.g. for attention modulation effect, either attended-unattended or unattended-attended was used), then the maximum size of significant cluster was recorded for this permutation. The procedure was repeated 10,000 times to get the null distribution, and the p value can be derived for a given cluster size.

## Results

### GE BOLD showed additive effect of attention on contrast responses, stronger in the superficial and deep cortical depths of early visual cortex

In experiment 1, we investigated the effect of spatial attention on contrast response functions of T2* BOLD signals across cortical depths in human early visual cortex. If attention operate through feedback visual pathways, we expect to see strongest attention modulation in the superficial and/or deep layers of early visual cortex. Otherwise, if attention modulate early visual activity through feedforward connections from subcortical pathway, attention modulation should be strongest in the middle layers. During the experiment, subjects were required to maintain fixation and to pay attention to the cued checkerboard to detect occasional spatial frequency change of the stimulus (Figure 2). Stimuli were presented at five contrast levels. In a pilot experiment, task difficulties were matched across different contrast conditions with a staircase procedure. Subjects’ performance in the scanner showed no significant difference across different contrast conditions (2.5%: 81.4±3.4%, 6.0%: 85.5±3.8%, 14.4%: 83.8±4.2%, 34.6%: 88.3±3.7%, 82.9%: 87.6±2.5%, F(4,14)=0.63, p=0.644).

To investigate the gain control mechanism of attention associated with feedforward or feedback connections, the Naka-Rushton equation (Eq. 1) was used to fit the BOLD contrast response functions (CRFs) across cortical depth in the early visual areas (Naka and Rushton 1966, Boynton, Engel et al. 1996). Independent variable c is the luminance contrast of checkerboard pattern, C_50_ is the contrast at which the response reaches half of its maximum dynamic range thus indicates contrast sensitivity, R_max_ is the maximum response from baseline, and parameter b denotes the baseline activity.

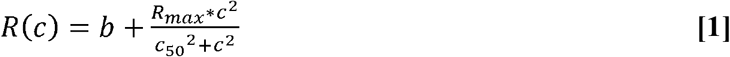

To understand the mechanism of attention modulation on BOLD CRFs, seven models based on Naka-Rushton equation were fitted to the group averaged contrast response functions: a full model with all parameters free to change, and six reduced models including baseline shift, response gain, contrast gain, response gain + baseline shift, contrast gain + baseline shift, and response gain + contrast gain. Data from the attended and unattended CRFs were fitted together. For the unattended CRF model, parameter b was fixed at zero. An F-test was used to compare the goodness of fit between different models (Li, Lu et al. 2008). Figure S3 shows the variance explained from different models (only 5 models with most variance explained were shown). Among all reduced models, baseline shift + contrast gain explained the most variance across ROIs. However, it showed no significant difference with the baseline shift model which has the fewest model parameters (p>0.8 for all ROIs). Furthermore, we used Akaike Information Criterion corrected for small sample size (AICc) for model comparison (Anderson and Burnham 2004), the baseline shift model gave the smallest AICc among all models tested. Thus we selected the baseline shift model as the best model that explained the most variance of data with the least number of parameters. Figure 3A shows the group averaged contrast responses and fitted curves of the baseline shift model. Such additive shift of attention on BOLD CRF is consistent with previous BOLD fMRI studies of attention on contrast responses (Buracas and Boynton 2007, Li, Lu et al. 2008, Pestilli, Carrasco et al. 2011).

**Figure 3.**
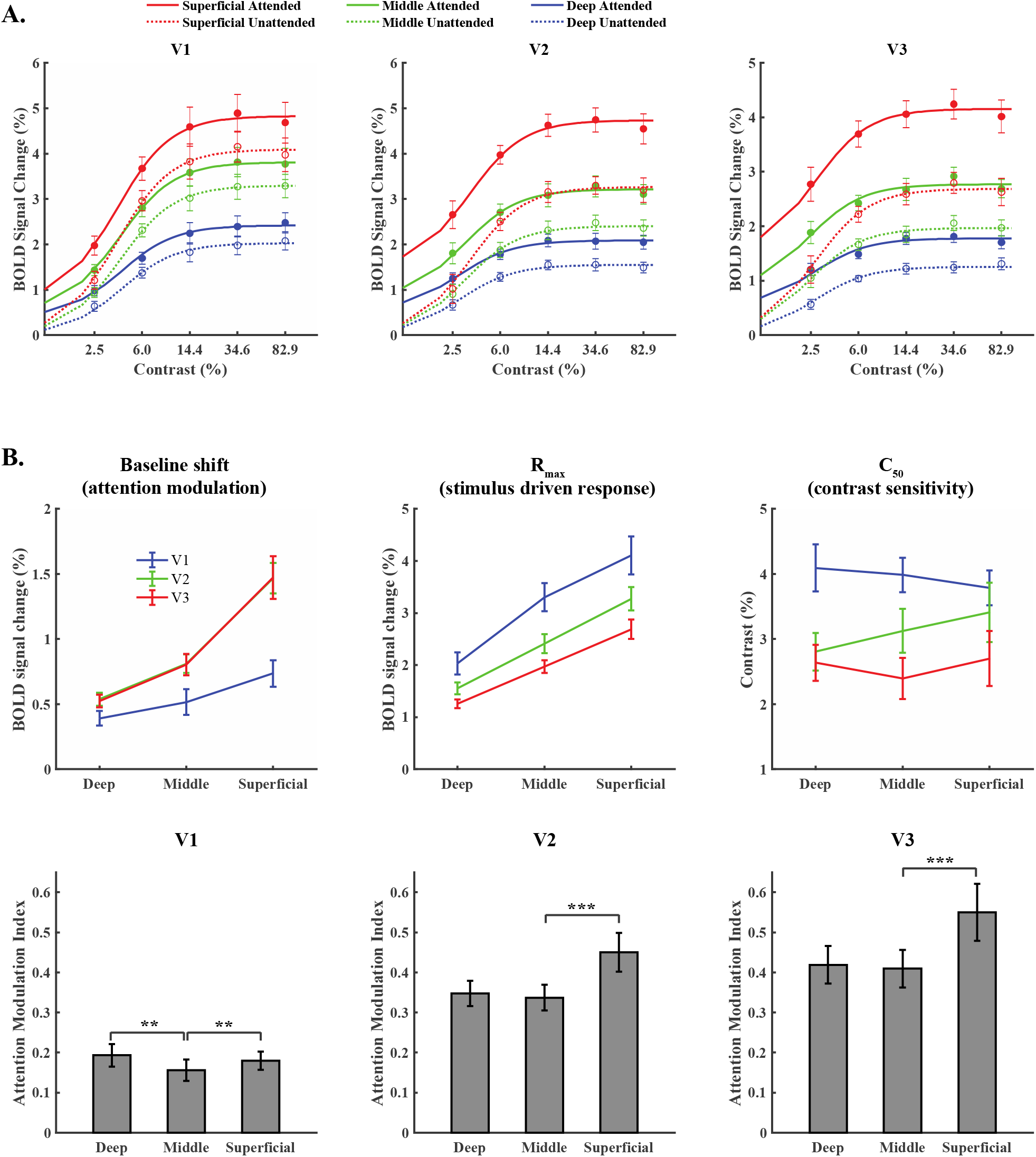
**A. BOLD contrast response functions from different cortical depths in V1, V2 and V3.** Solid and dotted curves represent the fitted responses from the baseline shift model of Naka-Rushton equation. Error bars represent standard deviation of bootstrapped distribution. **B. Upper row:** Parameters of the baseline shift Naka-Rushton model of attention across cortical depths. **Lower row:** Attention modulation index across cortical depths of V1-V3. Error bars represent standard deviation of bootstrapped distribution of the mean. ** and *** indicate p<0.01 and p<0.001 after multiple comparisons correction.

Figure 3B (upper panel) shows the mean and standard error of the baseline shift model parameters across cortical depth in V1-V3. The baseline shift effect of attention (parameter b) was stronger in V2 and V3 compared to V1 (both p<0.001), and increased monotonically from deep to superficial depth. The maximum stimulus driven response from baseline (R_max_) decreased from V1 to V3 (V1 vs. V2, p=0.011, V2 vs. V3, p=0.016), and also showed a monotonic increase toward the superficial depth. C_50_ decreased from V1 to V3 (V1 vs. V2, p=0.029; V2 vs. V3, p=0.12, V1 vs. V3, p<0.001), indicating an increase of contrast sensitivity along the ascending visual pathway. However, contrast sensitivity (C_50_) showed no significant difference across cortical depth (all p>0.05).

The monotonic increase of attention modulation and stimulus driven response from deep to superficial cortical depth can be explained by the draining vein effect, which is the drainage of deoxygenated hemoglobin along ascending veins, causing stronger BOLD signals toward the cortical surface (Zhao, Wang et al. 2004, Uludag and Blinder 2017). This effect was consistent with previous cortical depth-dependent BOLD fMRI studies using gradient echo sequences (Polimeni, Fischl et al. 2010, Marquardt, Schneider et al. 2018). To minimize the influence from draining veins which is a scaling effect (Uludag and Blinder 2017, Marquardt, Schneider et al. 2018), we calculated a normalized attention modulation index by dividing the baseline shift effect of attention (parameter b) by the maximum stimulus driven response (R_max_)(O’Connor, Fukui et al. 2002, Schneider and Kastner 2009). It denotes the ratio between attention modulation and bottom-up stimulus driven responses. Figure 3B (lower panel) shows the results for this normalized index of attention modulation across cortical depth in different visual areas. Compared to the middle depth of V1, the attention modulation index was significantly larger in the superficial and deep cortical depths (superficial vs. middle, p=0.003, deep vs. middle, p=0.009, after Bonferroni correction, number of comparisons k=2). In V2 and V3, attention modulation index was significantly larger in the superficial than in the middle cortical depth (V2, p<0.001, V3, p<0.001, after Bonferroni correction). These findings were consistent with the hypothesis that top-down spatial attention modulates neural responses through feedback connections along the descending visual pathway.

### Ultrahigh resolution bSSFP fMRI revealed multiplicative effect of attention in V1 superficial and deep layers

Human primary visual cortex is about 2mm thick. Even with 0.75mm isotropic resolution in the GE-EPI experiment, there are only 2-3 voxels sampling across the cortical layers of V1. Image blur due to T2* decay and long EPI readout further reduced the effective spatial resolution. Thus to better reveal the laminar profile of attention modulation, we used ultra-high resolution bSSFP fMRI to investigate the effect of spatial attention on the contrast responses in human primary visual cortex V1. As shown by figure 4A (left panel), a single 3mm thick slice with 0.3mm in-plane resolution was placed perpendicular to a flat portion of calcarine sulcus in V1. Stimulus and procedure were similar to those in the GE-EPI experiment, except that visual stimuli were presented at low contrast (5%) and high contrast (50%) conditions. Task difficulty were matched with a staircase procedure in a pilot experiment. Subjects’ performance in the scanner showed no significant difference between the two contrast conditions (low contrast: 85.3±3.4%, high contrast: 88.5±1.5%, t(1,11) = 1.2, p = 0.259). Figure 4A (right) shows the activation map from a representative subject. Robust activations can be found in V1 gray matter, with relatively small signal change (~ 1.5% for high contrast stimulus). Pixels containing large draining veins showed large signal change (over 5%), consistent with strong intravascular contribution to bSSFP signals due to water diffusion in and around red blood cells (Dharmakumar, Hong et al. 2005). However, the intravascular activations were well contained within the blood vessels, without contaminating activations of surrounding gray matters. Pixels from large draining veins were later removed by a signal change threshold (Figure S4). Activation (beta) maps from figure 4A (right) shows that attention modulation was stronger in the high contrast condition than in the low contrast condition, indicating a multiplicative scaling effect (response or activity gain) of attention on contrast responses. Moreover, in the high contrast condition, strong attention modulation can be clearly seen in the deep cortical depth. Figure S5 shows the maps of attention modulation in the high contrast condition across the ROIs of all subjects, two layers of activation in the superficial and deep cortical depths can be seen for most subjects.

**Figure 4.**
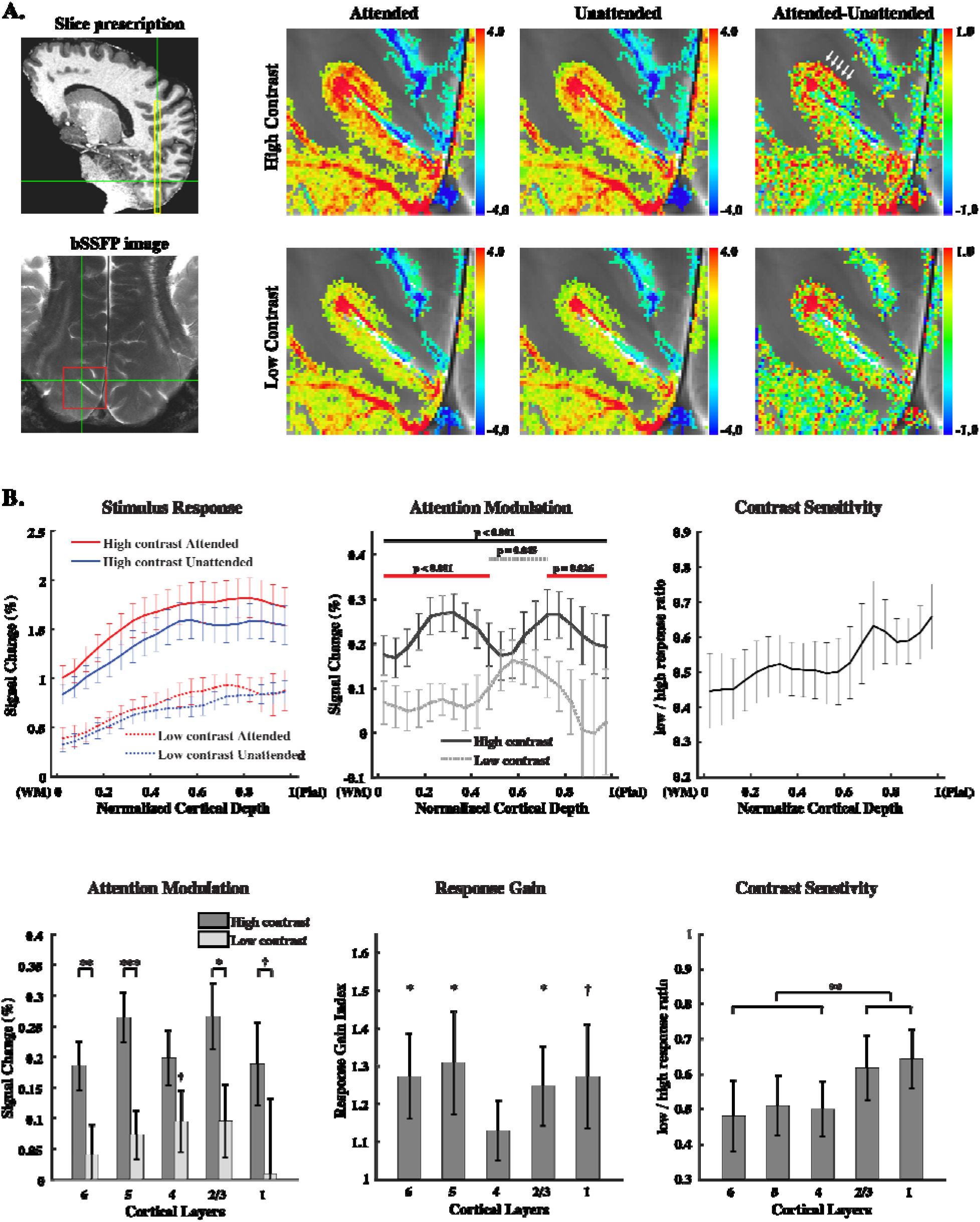
**A. Left:** Slice prescription and ultra-high resolution passband bSSFP image of a representative subject. A single bSSFP slice (3mm thick, 0.3mm in-plane resolution) was positioned perpendicular to a flat portion of calcarine sulcus on the left hemisphere. Yellow box indicates the slice position for the bSSFP image. Red square highlights the zoomed-in area in the right panels. **Right:** Activation maps from the same subject. Beta maps were thresholded at attended+unattended p<0.05 uncorrected. Clear deep layer attention modulation can be seen in the high contrast condition, as indicated by the white arrows. Color bars indicate percent signal change. **B. Upper row:** The laminar profiles of bSSFP fMRI responses to the visual stimuli (left), attention modulation (middle) and contrast sensitivity index (right). Horizontal bars in the middle panel indicate significant cortical depths after multiple comparison correction by cluster-based permutation test. Solid Black bar: significant attention modulation in high contrast condition; Dotted gray bar: significant attention modulation in the low contrast condition; Solid red bar: Significantly stronger attention modulation in the high contrast condition than in low contrast condition. **Lower row:** Attention modulation, response gain and contrast sensitivity in cortical depths corresponding to the locations of cortical layers. *, ** and *** indicate p < 0.05, p < 0.01, p < 0.001, respectively after multiple comparisons correction (Bonferroni). † indicates p < 0.05 before correction. Error bars represent standard deviation of bootstrapped distribution of the mean.

Group averaged laminar profiles of bSSFP fMRI responses to the visual stimuli show a monotonic increase toward the cortical surface (Figure 4B, upper row, left panel), but are relatively flat compared to the depth-dependent response of GE BOLD (Figure 3B). Attention modulation was stronger in the high contrast condition compared with the low contrast condition (Figure 4B, upper row, middle panel), indicating response or activity gain of attention. Moreover, such multiplicative scaling effect of attention were significant in the superficial and deep cortical depths (p<0.001 and p=0.026, respectively, after multiple comparison correction by cluster-based permutation test, see statistics in method section for details), but not in the middle cortical depths. This finding was consistent with the V1 results from the GE-EPI experiment.

The laminar profiles of attention modulation show clear distinction in the high contrast and low contrast conditions. In the high contrast condition, attention modulation was significant across all cortical depths but a clear double-peak signature can be seen in the superficial and deep layers; whereas in the low contrast condition, attention modulation was significant only in the middle cortical depths (p=0.045 after correction). To characterize the differences in contrast sensitivity across cortical depths, a contrast sensitivity index was calculated as the ratio between the low contrast and high contrast responses in the unattended condition. The laminar profile shows higher contrast sensitivity in the superficial cortical depths than those in the middle and deep cortical depths (Figure 4B, upper row, right panel).

In lower panels of Figure 4B, the laminar profiles were resampled into five depth bins according to the thickness of cortical layers of human primary visual cortex (Burkhalter and Bernardo 1989, de Sousa, Sherwood et al. 2010, Amunts, Lepage et al. 2013, Balaram, Young et al. 2014, Wagstyl, Larocque et al. 2019): layer 6, 20%, layer 5, 10%, layer 4, 35%, layer 2/3, 25%, layer 1, 10% (values rounded to the nearest multiple of 5). As shown by the left panel (Figure 4B, lower row), attention modulation in the high contrast condition was stronger than in the low contrast condition, which was significant in layer 2/3, layer 5, layer 6 (p=0.0135, 0.0005 and 0.0045, respectively, Bonferroni corrected, number of comparisons k=5) and layer 1 (p=0.0420, uncorrected), but not in layer 4 (p=0.0638, uncorrected). In the low contrast condition, attention modulation was only significant in layer 4 (p=0.0283, uncorrected). To further quantify the response or activity gain of attention, a response gain index was calculated by Eq. [2].

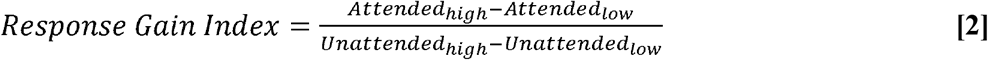

As shown by the middle panel (Figure 4B, lower row), significant response gain can be found in layer 6, layer 5, layer 2/3 (p=0.0319, 0.0455 and 0.0318, respectively, Bonferroni corrected) and layer 1 (p=0.0205, uncorrected), but not in layer 4 (p=0.0601, uncorrected). The average response of superficial and deep layers (layer 1, 2/3, 5,6 combined) showed significantly stronger response gain than layer 4 (p=0.0221). Contrast sensitivity was significantly stronger in the superficial layers (layer 1/2/3) than in the middle and deep layers (layer 4/5/6, p=0.0070), consistent with thalamocortical projections mainly targeting layer 4 and layer 6 (Hubel and Wiesel 1972, Hendrickson, Wilson et al. 1978, Thomson 2010), and relatively linear contrast responses in the lateral geniculate nucleus of the thalamus (Kastner, O’Connor et al. 2004).

In the GE-EPI experiment, we found an additive effect of attention on the contrast responses of early visual cortex, while passband bSSFP fMRI showed multiplicative scaling effect of attention in V1 gray matter. It is generally considered that T2* weighted GE BOLD at ultrahigh field are strongly contributed by extravascular macrovessel activity, such as large draining veins (Kim 2018); while bSSFP fMRI at short TR mainly captures diffusion activity around small vessels in the gray matter (Lee, Dumoulin et al. 2008, Miller 2012, Scheffler, Heule et al. 2018). Thus we hypothesize that the discrepancy from the two pulse sequences might be related to the difference of attention modulation on the responses from large draining veins and microvessel activity from gray matter. To further test this hypothesis, we examined the attention modulation effect on the bSSFP signals from V1 gray matter and large draining veins. From the bSSFP activation map, extravascular signals in gray matter and intravascular signals from large draining veins can be clearly distinguished (Figure 4 and Figure S4). As shown by Figure 5 (right panel), attention modulation on the bSSFP fMRI signals from large draining veins was additive (or non-multiplicative): the effect size of attention modulation was comparable between low contrast and high contrast conditions. Two-way ANOVA showed a significant main effect of attention (F(1,11)=45.7, p<0.001) and contrast (F(1,11)=21.9, p<0.001), without significant interaction between attention and contrast (F(1,11)=1.2, p=0.3). However, for microvascular response in the gray matter, there was a significant interaction between attention and contrast (F(1,11)=15.1, p=0.003), indicating stronger attention modulation in the high contrast condition than in the low contrast condition. These findings clearly demonstrated that attention modulation on the responses of large draining veins was additive shift, consistent with previous attention studies using T2* BOLD fMRI at 3T (Buracas and Boynton 2007, Li, Lu et al. 2008, Pestilli, Carrasco et al. 2011). However, attention modulation was multiplicative (response or activity gain) on extravascular microvessel activity in gray matter, consistent with the findings from electrophysiology (Di Russo, Spinelli et al. 2001, Reynolds and Heeger 2009).

**Figure 5.**
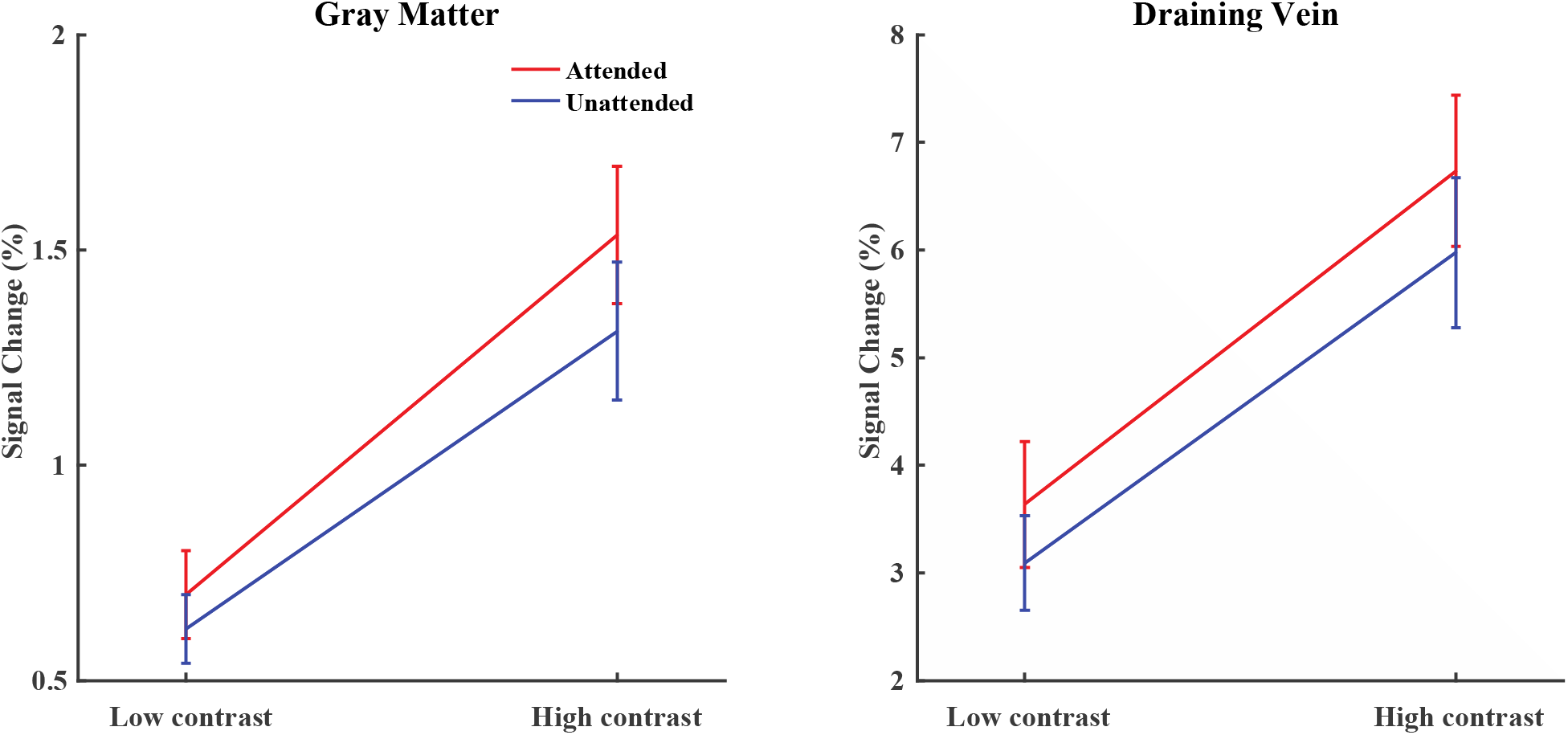
bSSFP fMRI responses from gray matter and large draining veins. **Left.** Attention modulation of contrast responses shows multiplicative scaling effect in V1 gray matter: stronger attention modulation in the high contrast condition than in the low contrast condition. **Right.** Attention modulation of contrast responses from large draining veins shows additive effect: similar attention modulation between low contrast and high contrast conditions. Error bars represent standard error of the mean.

### Stronger nonlinearity of BOLD signal from large than small blood vessels could explain the additive vs. scaling effect of attention

Our experiments clearly demonstrate that T2* weighted GE BOLD signals dominated by large vessels showed additive (or non-multiplicative) effect of attention on contrast responses, while T2 weighted fMRI signals from microvessels revealed multiplicative scaling effect of attention. Intuitively, microvessels are closer to the site of neural activity, thus it seems reasonable that high fidelity T2 weighted responses from microvessels showed similar results as neurophysiological recordings. Recently, an elegant study from Bao et.al demonstrated a strong nonlinear relationship between T2* BOLD signals and neural activity using achiasma human visual system (Bao, Purington et al. 2016). Thus we hypothesize that the additive shift of attention on T2* BOLD signal might be due to the strong nonlinear relationship between BOLD signal of large blood vessels and underlying neural activity, while the responses from microvessels are relatively linear thus preserved the multiplicative scaling effect of neural modulations. To test this hypothesis, we used a BOLD biophysics model (Davis model) to simulate the nonlinearity of BOLD signals from large and small blood vessels as a function of cerebral metabolic rate for oxygen (CMR_O2_), which directly reflects the energy requirement of underlying neural activity. Eq. [3] shows the nonlinear model describing the relationships between BOLD signal change and oxygen consumption and vascular responses (Davis, Kwong et al. 1998).

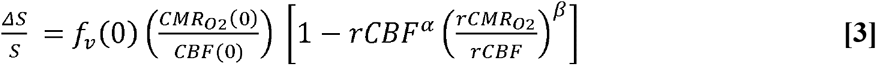

*f*_*ν*_ is the fraction of blood volume. rCBF and rCMR_O2_ are normalized ratio of CBF and CMR_O2_ relative to baseline. α denotes the exponent representing the nonlinear relationship between CBV and CBF. For large vessels we set *α* = 0.28 (Chen and Pike 2009), for small vessels *α* = 0.1 (Stefanovic, Hutchinson et al. 2008). The exponent *β* models the influence of deoxyhemoglobin on transverse relaxation which is assumed to be smaller for larger vessels and at higher magnetic fields. Here we set *β* = 0.8 for large vessels, and *β* = 1.2 for small vessels (Martindale, Kennerley et al. 2008, Griffeth and Buxton 2011). The coupling ratio between CBF and CMR_O2_ change was set at 2 (Stefanovic, Warnking et al. 2004). Indeed, the simulation result (Figure 6) showed strong nonlinearity of BOLD signals for large blood vessels, which transforms the scaling effect of attention into an “additive offset”. BOLD signal change of small vessels is relatively linear, which largely preserves the scaling effect of attention modulation.

**Figure 6.**
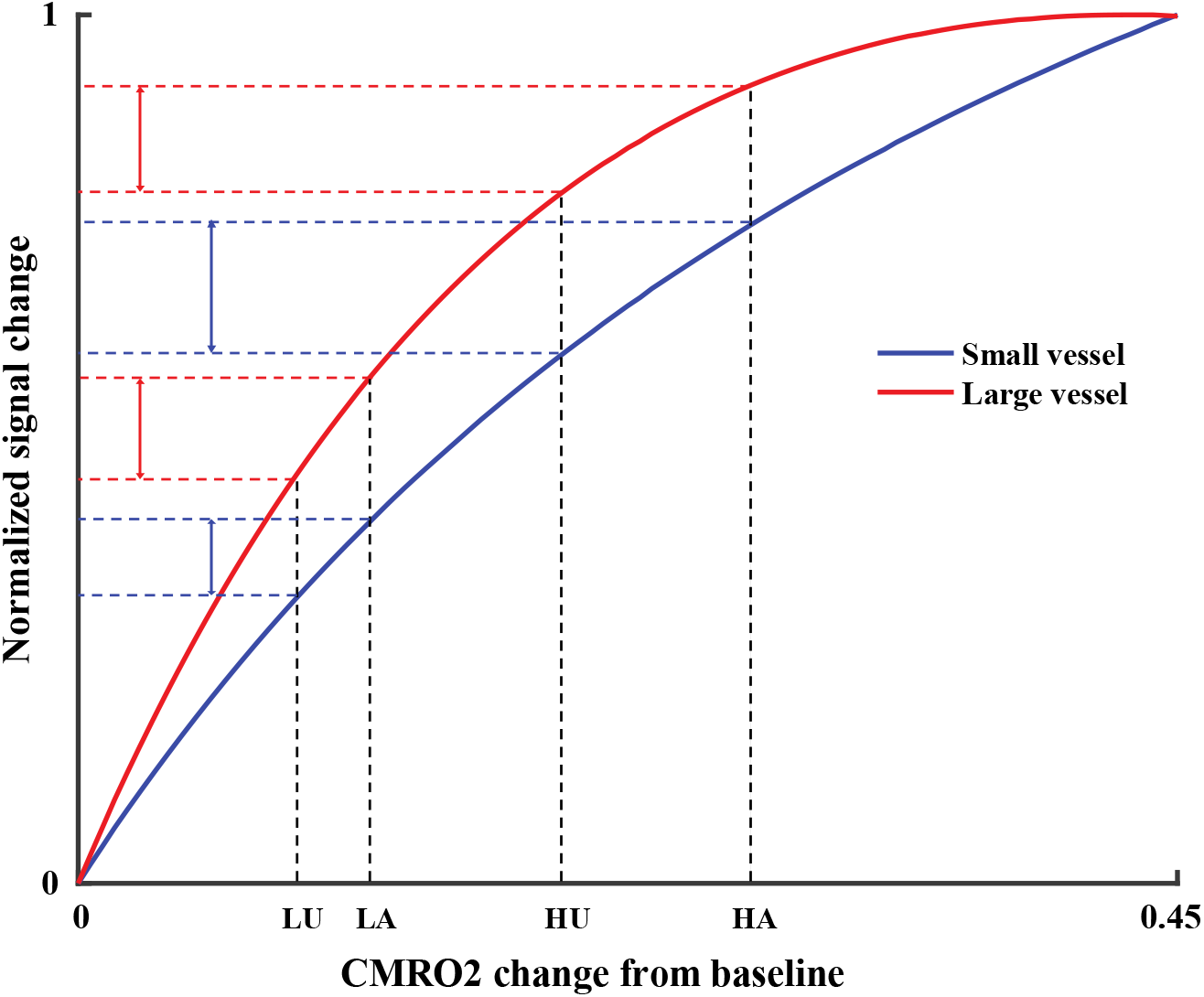
Difference in BOLD nonlinearity between large and small blood vessels explains the additive and scaling effect of attention. **LU**: Low contrast stimulus, Unattended, **LA**: Low contrast stimulus, Attended, **HU**: How contrast stimulus, Unattended, **HU**: High contrast stimulus, Attended.

## Discussion

### Top-down spatial attention mainly modulates early visual activity through feedback connections

Using submillimeter resolution fMRI at 7T, we investigated cortical depth-dependent attention modulation of contrast responses in human early visual cortex. In experiment 1, gradient echo T2* weighted BOLD signal showed additive offset of attention on contrast responses across different cortical depths of V1, V2 and V3. Relative to the stimulus driven response, attention modulation was stronger in the superficial and deep cortical depths than in the middle depth of V1, consistent with the hypothesis that attention modulates early visual activity through direct cortico-cortical feedback connections. In V2 and V3, attention modulation were strongest in the superficial cortical depth, supporting cortico-pulvino-cortical feedback to the superficial layers of extrastriate visual cortex (Shipp 2003). Consistent with previous studies (O’Connor, Fukui et al. 2002), we also found stronger attention modulation in V2 and V3 compared to V1. These findings support the idea that attention operates through feedback connections hierarchically along the descending visual pathway. In experiment 2, using ultra-high resolution (0.3mm in-plane) distortion free T2 weighted bSSFP fMRI method, we found response or activity gain of attention on V1 contrast responses: attention modulation was stronger in the high contrast condition than in the low contrast condition. This multiplicative effect of attention was significant in V1 superficial and deep layers, but not in the middle layer. This finding further supports that attention modulates V1 activity through direct cortico-cortical feedback.

Therefore, both experiments consistently showed that top-down spatial attention can modulate contrast responses in the superficial and deep layers of human early visual cortex, supporting the hypothesis that attention operates through feedback connections along descending visual pathway in a hierarchical fashion. These findings are consistent with recent monkey electrophysiology studies showing that top-down attention enhanced multi-unit activity mainly in the superficial and deep layers of V1 (van Kerkoerle, Self et al. 2017), reduced correlated neural activity in the superficial layers of V1 (Denfield, Ecker et al. 2018), and reduced the variability of neural activity in the superficial layer of V4 (Nandy, Nassi et al. 2017).

### A distinct mechanism of attention is associated with feedforward input in V1 middle layer

Interestingly, our bSSFP results showed a distinct laminar profile of attention modulation in the low contrast condition compare to the high contrast condition. Attention modulation in the low contrast condition was strongest in the middle depths of V1 and showed no significant difference compared with the high contrast condition, indicating baseline enhancement or contrast gain of attention modulation in V1 middle layers. This finding suggests a distinct mechanism of attention associated with feedforward visual activity through subcortical neural circuits (O’Connor, Fukui et al. 2002, McAlonan, Cavanaugh et al. 2008). Attention increased contrast sensitivity in V1 middle layer is also consistent with a previous study showing that visual perceptual learning increased the contrast sensitivity of Magnocellular response of the human LGN (Yu, Zhang et al. 2016). A recent monkey electrophysiology study also found different neural mechanisms of attention modulation in the superficial and middle layers of V4 (Nandy, Nassi et al. 2017). Altogether, our bSSFP fMRI results highlight an important point that spatial attention could modulate early visual activity through distinct feedback and feedforward visual pathways, with different gain control mechanisms of attention.

A recent lamina-resolution fMRI study using T2* BOLD showed that compared to the middle layer, feature-based attention modulation was stronger in the superficial layer of V1 and V3, but not in V2 (Lawrence, Norris et al. 2019). However, they didn’t show strongest attention modulation in the middle layer in the low contrast condition. Such inconsistency between the previous study and our study might be due to the difference of feature-based attention and spatial attention, and difference in fMRI contrast mechanisms.

### Nonlinearity of BOLD signals from large draining vein causes the additive effect of attention on contrast response functions

Across cortical depths, attention modulation of GE T2* BOLD signals showed additive offset on contrast response function, without significant difference of attention modulation between different contrast conditions. This finding was consistent with previous 3T BOLD fMRI studies that mainly revealed additive effect of attention modulation on contrast response functions in human visual cortex (Murray and He 2006, Buracas and Boynton 2007, Li, Lu et al. 2008, Pestilli, Carrasco et al. 2011). However, neurophysiological studies usually found contrast gain or response gain of attention modulation on contrast responses (McAdams and Maunsell 1999, Reynolds, Pasternak et al. 2000). There are a number of potential explanations to explain such discrepancy of attention modulation between BOLD fMRI and neural activity. One important consideration is that fMRI signals reflect pooled responses from large populations of neurons. For example, attention might increase baseline firing rate of large unresponsive neural populations, which might dominate fMRI response but is negligible from single unit recordings. Attention modulation on pooled neuronal activity with response gain and contrast gain could also behave like an additive effect (Williford and Maunsell 2006, Hara, Pestilli et al. 2014). However, our bSSFP fMRI data also reflected population response but clearly showed multiplicative scaling effect of attention. Moreover, human EEG studies found response gain of attention modulation on contrast responses at the population level (Di Russo, Spinelli et al. 2001, Kim, Grabowecky et al. 2007). Another possibility is that BOLD signals reflected subthreshold activity which is more related to LFP rather than spike rate, and attention might influence the subthreshold response differently from spiking activity. But this cannot explain why BOLD signal showed additive offset but EEG activity showed scaling effect. Finally, it has been suggested that attention might influence vascular responses differently from neural activity (Buracas and Boynton 2007). However, there is no clear evidence to support this hypothesis.

In the current study, we found that the additive effect of attention on contrast responses were associated with BOLD signals from large draining veins but not with microvessel activity in the gray matter, and that such differences can be explained by the nonlinear relationship between BOLD signal and vascular response. At both 3T (previous studies) and 7T (the current study), GE T2* BOLD signals are highly sensitive to the susceptibility effect of large blood vessels, and both showed additive offset of attention modulation. At 7T, T2 weighted bSSFP BOLD signal in gray matter mainly reflected extravascular diffusion activity around small vessels, and revealed multiplicative scaling effect of attention. In the bSSFP experiment, intravascular response from large draining veins also showed additive effect of attention on contrast responses. In a simulation analysis, we found that compared to microvessels, BOLD signal from large blood vessel exhibits stronger nonlinearity, which could transform a multiplicative scaling effect of attention modulation into an additive effect. Therefore, these evidence suggest that the strong nonlinearity of BOLD signal from large draining veins could cause the additive effect of attention on contrast response functions. An important implication of these findings is that compared to traditional GE T2* BOLD, fMRI signals with higher tissue specificity such as T2 BOLD, CBV and CBF, may have a more linear relationship with underlying neural activity. Future studies should investigate the effect of attention modulation using calibrated fMRI (CMR_O2_), or other quantitative fMRI methods, such as CBF- and CBV-based sequences.

### Advantage and disadvantage of bSSFP imaging for high resolution fMRI at ultrahigh field

The current study demonstrates that balanced SSFP can be a useful method for high resolution fMRI of cortical layer-dependent activity in the human brain. The laminar profile of bSSFP fMRI signals (Figure 6A) showed weaker bias toward the cortical surface compared to the response profile of GE BOLD signals (Figure 4A, middle panel). This finding is consistent with previous layer-fMRI studies using SE sequences (Zhao, Wang et al. 2006, Olman, Harel et al. 2012, De Martino, Zimmermann et al. 2013), thus supporting the theory and simulation that bSSFP fMRI signal at short TR primarily detects diffusion activity around small vessels (Scheffler, Heule et al. 2018). Passband bSSFP fMRI signals also helped to reveal the neural effect of attention modulation, suggesting a closer relationship of T2 BOLD with underlying neural activity. A major advantage of bSSFP imaging is the minimal spatial distortion or blurring due to extremely short readout time. Distortion free is important for registration with anatomical images and accurate segmentation of cortical surfaces, reduced blurring is also critical for high resolution studies. Our bSSFP data showed strong intravascular activity in large draining veins, but they are well contained in the blood vessel without contaminating activation in the gray matter. Large pial veins with strong signal change can be easily removed by post-hoc data analysis approach, but smaller penetrating vessels are hard to remove and will reduce the spatial specificity of bSSFP signals. A major disadvantage of bSSFP is the limited brain coverage due to much reduced acquisition speed compared to EPI-based methods. In the current study, motion correction for a single image slice was quite challenging thus we sacrificed higher order visual areas V2 and V3 because of limited coverage. Future studies should combine different acceleration techniques, such as parallel imaging, partial Fourier acquisition, multiline and parallel transmission to achieve high speed 3D-acquisition.

## Conclusions

Top-down spatial attention showed additive offset on T2* weighted GE BOLD but multiplicative scaling on T2 weighted bSSFP fMRI signals in the superficial and deep layers of human early visual cortex. Attention modulation of low contrast response of T2 BOLD was also strongest in the middle cortical depths of V1. These findings demonstrate that top-down spatial attention mainly operates through feedback connections along descending visual pathways, and a distinct mechanism of attention may also be associated with low contrast feedforward input through subcortical pathway. Finally, the neural mechanism of attention modulation in the agranular layers should be response or activity gain, because the additive effect of attention on T2* BOLD can be explained by the nonlinearity of BOLD signals from large blood vessels.

## Supplemental Materials

**Figure S1.**
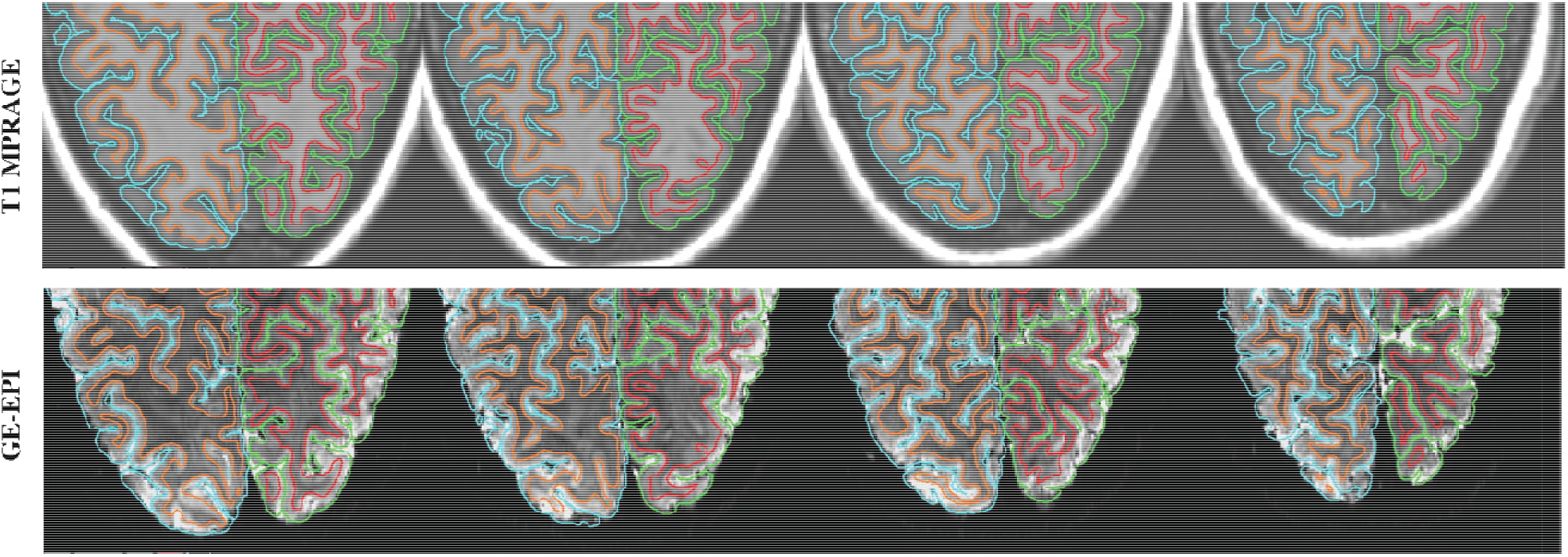
Illustration for the quality of surface segmentation and image registration. A representative subject from the GE-EPI experiment was shown here.

**Figure S2.**
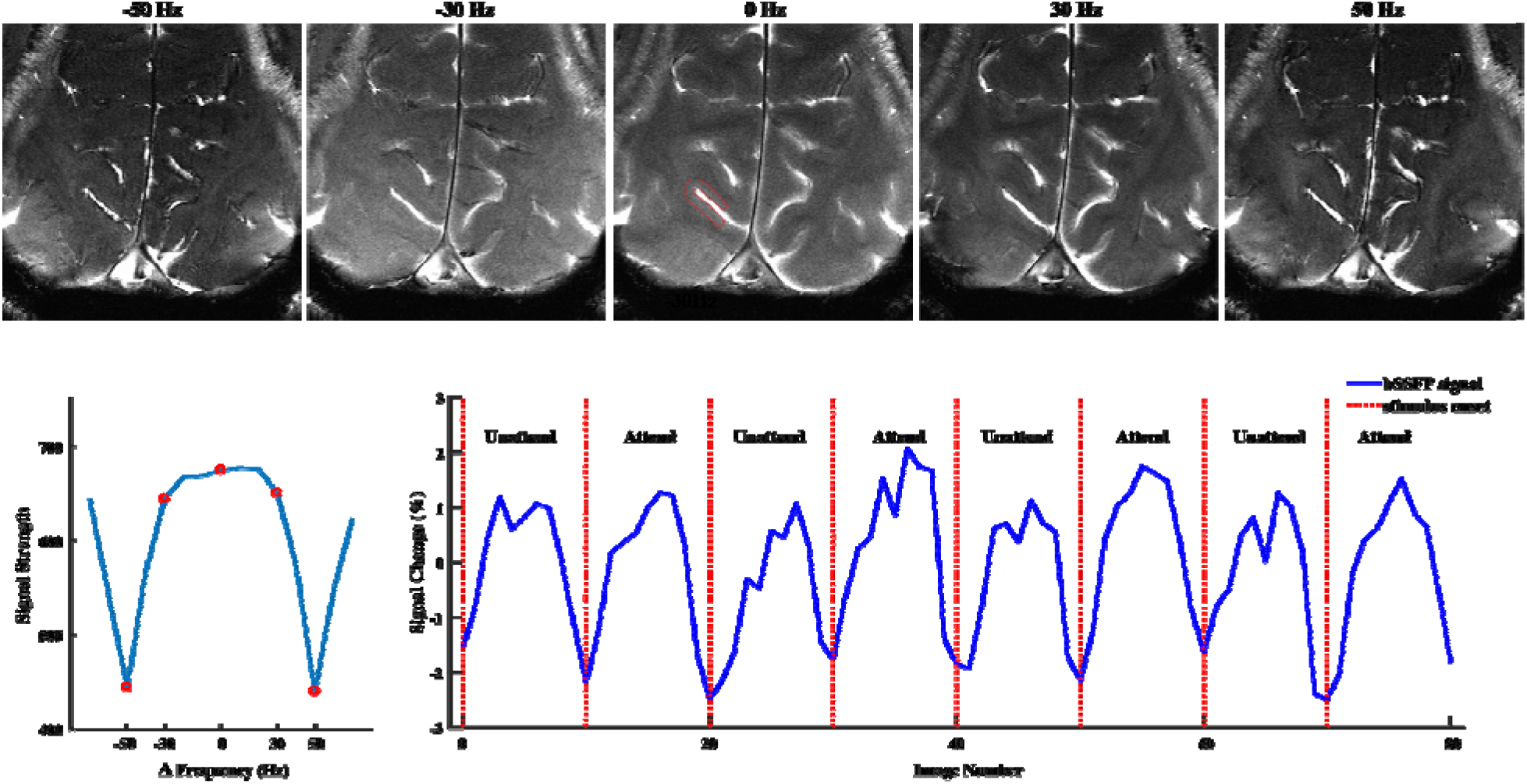
Selection of delta frequency of RF pulse for the bSSFP acquisition. **Upper row:** A series of bSSFP images were collected using RF pulses with a parametric offset from the on-resonance frequency (−70~70 Hz in a step of 10 Hz, only five images were shown here). **Lower left:** Off resonance profile from the selected ROI (as indicated by the red frame from above), red circles correspond to the images from upper row. For this representative subject, RF delta frequency of 0 Hz was located in the middle of the passband, thus was used for functional data acquisition. **Lower right:** ROI time course from a single run shows good signal to noise ratio (high contrast stimulus). The difference between attended and unattended condition can be clearly seen.

**Figure S3.**
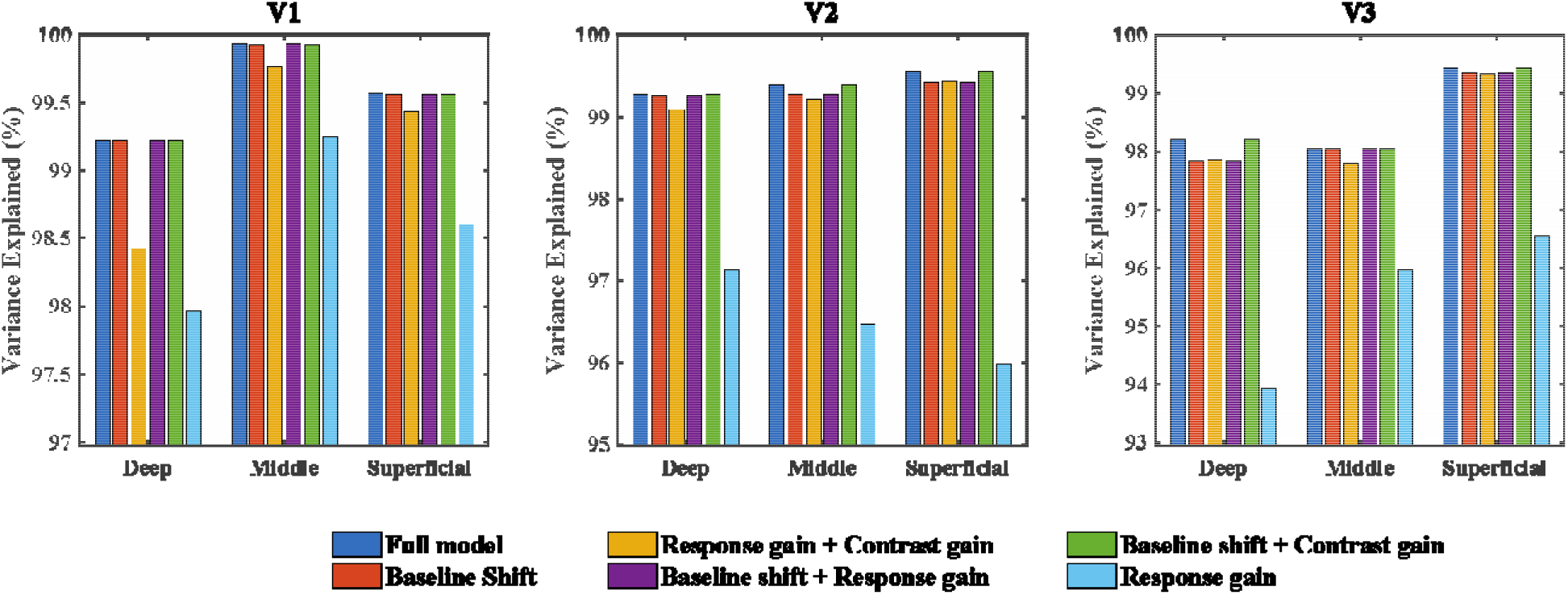
shows the explained variance of BOLD contrast response functions in the GE-EPI experiment.

**Figure S4.**
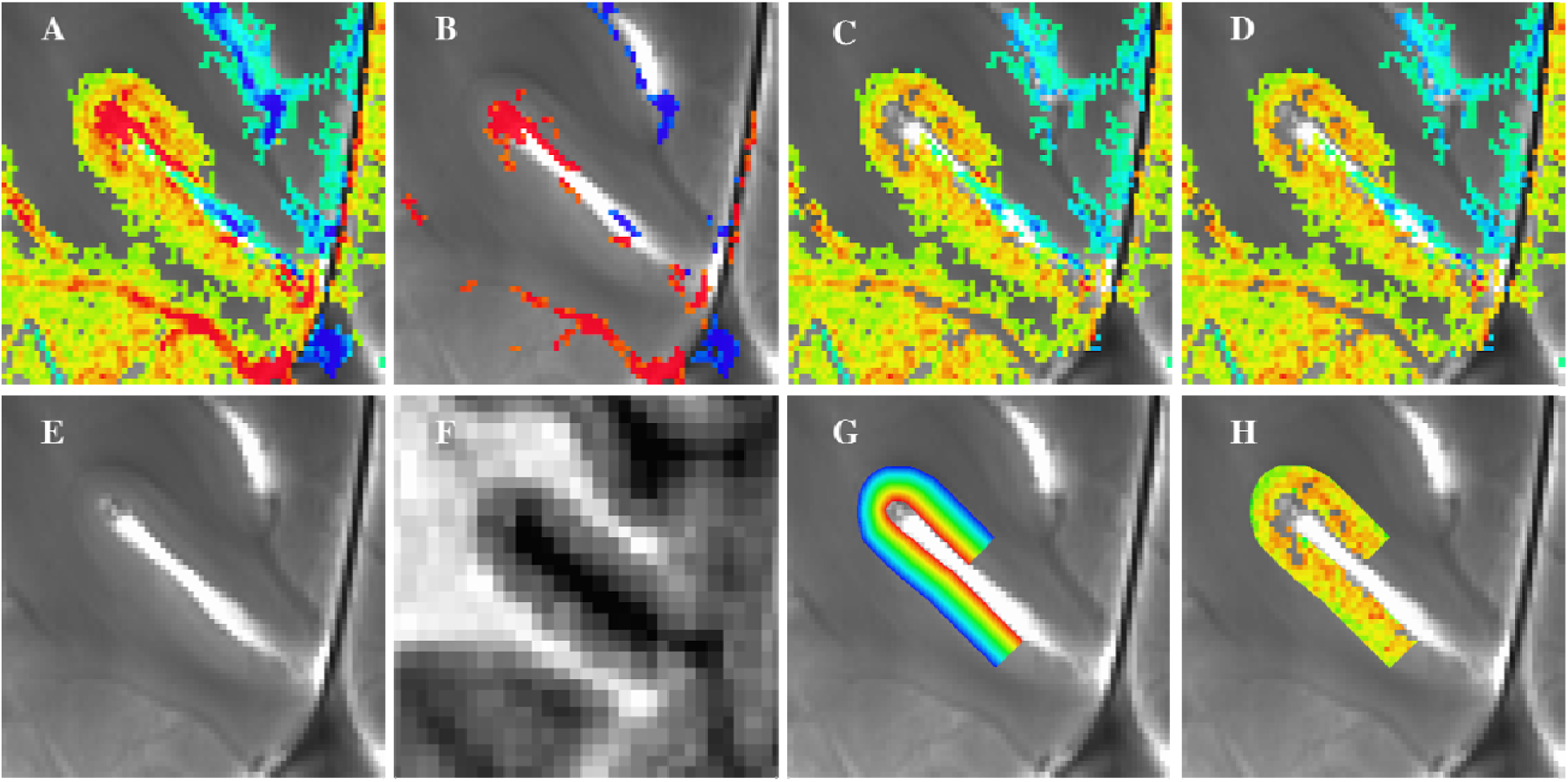
Vein removal and ROI definition in the bSSFP experiment. **A.** Activation map across all conditions (thresholded at p<0.05). **B.** Activation from large draining veins. Only pixels with over 7% signal change were shown. **C.** Pixels from large draining veins in B were removed from A. **D.** Activation map in C was visually inspected, (two) remaining pixels from large pial veins were manually removed. **E.** High resolution bSSFP image provides good contrast between CSF and gray matter, and was used to define the gray matter (pial) surface. **F.** T1 MP2RAGE image shows good contrast between gray matter and white matter and was primarily used to define the WM surface. Definition of WM surface also referred to the bSSFP image and activation map. **G.** Manually defined ROI. Pixel color represents normalized cortical depth (Blue: WM surface, Red: pial surface). **H.** Activation (beta) map within the ROI. Pixels with large signal change (draining veins) were removed.

**Figure S5.**
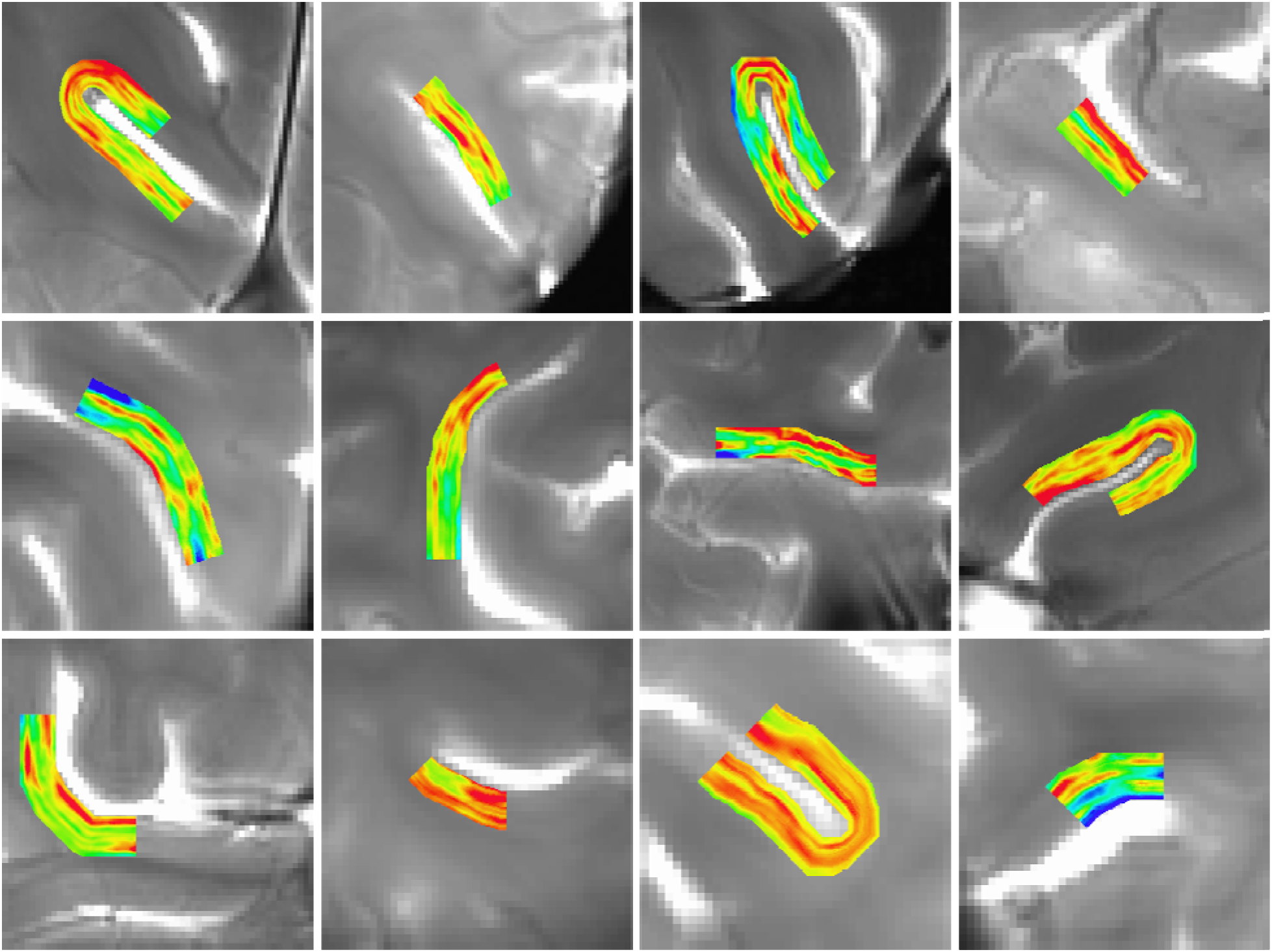
Pattern of attention modulation (attended-unattended) in the high contrast condition. Each figure shows the result from one subject. For illustration purpose, pattern was smoothed within-layer with a 1mm FWHM Gaussian window. Double-layered pattern in the superficial and deep cortical depths can be seen for most subjects.

